# A host-adapted commensal fungus from pet store mice drives type 2 immunity and cross-kingdom protection

**DOI:** 10.1101/2025.04.30.651489

**Authors:** Geongoo Han, Rebecca Yunker, Mohammad H. Hasan, Alex Bruce, Kellie Baldaro, Jessica Pacia, Nicholas N. Jarjour, Lalit K. Beura, Shipra Vaishnava

## Abstract

Naturalized, wilded, wildling, and dirty/pet store mouse models represent a spectrum of approaches designed to make laboratory mice more immunologically and physiologically similar to wild or human contexts by increasing their exposure to naturally occurring microbes and pathogens. In this study, we screened the gut mycobiome of pet store mice, and identified *Kazachstania pintolopesii* as a dominant fungus in pet store mice across various geographical locations. *K. pintolopesii* strains isolated from mice in geographically distinct pet stores stably colonize the gastrointestinal tract of laboratory mice, independent of gut bacterial composition, maintaining high fungal burdens for extended periods. *K. pintolopesii* rapidly became the dominant fungus in the mouse gut in conventional, antibiotic, and germ-free settings, outcompeting other non-murine fungal strains. Pet store-derived *K. pintolopesii* exhibited unique immunological properties distinct from typical anti-fungal responses. Unlike *C. albicans* colonization, *K. pintolopesii* did not induce circulating neutrophil expansion or Th17 cell populations in the gut mucosa. When administered systemically, *K. pintolopesii*-infected mice showed 100% survival with minimal fungal burden in kidneys, contrasting sharply with lethal *C. albicans* infections. Adaptive immune deficiency (Rag1 knockout mice) did not affect *K. pintolopesii* colonization or host response, indicating that B and T cell-mediated immunity does not restrain this fungus. *K. pintolopesii* colonization provided no cross-protection against systemic candidiasis further establishing lack of immune activation. These findings demonstrate that *K. pintolopesii* establishes a benign host-fungal relationship through neutrophil-independent mechanisms, avoiding classical anti-fungal immune activation while maintaining stable gut colonization. Instead, it selectively induces strong type 2 mucosal immune responses, increasing tuft and goblet cell counts and stimulating Th2 and group 2 innate lymphoid cell (ILC2) populations. This immune profile confers notable protection against intestinal nematode infection, demonstrated by reduced *Heligmosomoides polygyrus* egg counts. Altogether, *K. pintolopesii* serves as an exemplary model for commensal mycobiota, revealing distinct mechanisms for host tolerance and immune modulation.

## Introduction

Commensal microorganisms are crucial for host immunity and have coevolved with their natural hosts, often exhibiting high species restriction. This phenomenon is observed across various life forms. In invertebrates, the Hawaiian bobtail squid depends on *Vibrio fischeri* for tissue development ^1^, while tsetse flies rely on *Wigglesworthia glossinidia* for fitness and reproduction ^2^. Similarly, in mammals, gut immune cell maturation is driven by host-specific bacteria. Experiments have shown that human or rat gut bacteria introduced into germ-free (GF) mice are less effective at stimulating immune cell development compared to mouse-specific bacteria, highlighting the importance of host-adapted microbes in shaping host physiology ^3^. The mammalian gut houses a complex ecosystem of microbes, including bacteria, viruses, and fungi. However, our understanding of whether and how commensal fungi interact exclusively with their hosts is limited due to the lack of true commensal models that can stably colonize the mouse gut without causing inflammation or requiring immune suppression. Most research efforts have focused on *Candida albicans,* a gut commensal that is the main cause of opportunistic infections in humans. However, studying *C. albicans* commensalism in humans is somewhat limited because of technical and ethical challenges. As such introducing *C. albicans* to the murine gut has emerged as a dominant model system. But this approach often requires antibiotic treatment to maintain stable colonization and can induce inflammatory responses compromising interpretation ^4–6^. Therefore, alternative commensal fungi that have coevolved with their natural host are needed to better understand fungal-host symbiosis.

In recent years, researchers have embraced alternative models of microbial exposure for laboratory mice to better simulate the complex immune challenges faced by wild mammals. Approaches such as "re-wilded" models—where laboratory mice are released into natural environments^7^ or “wildling models” where embryos of laboratory mice are transferred into wild-caught female mice^8^—have been developed to introduce a broader range of natural microbiota and pathogens during early life. Another widely used strategy is the creation of "dirty mice," which involves co-housing standard laboratory mice with pet store or wild-derived mice, thereby facilitating the transmission of diverse microbes between animals and mimicking the microbial exposures found in free-living populations^9–11^. Studies using these models have demonstrated that increased microbial exposure leads to the development of an "immunologically experienced" immune system, marked by higher levels of memory T cells, diverse subsets of specialized immune cells, and enhanced inflammatory responses. The differences in microbial exposure between laboratory, wild, and pet store mice present a unique opportunity to identify and characterize potential commensal fungi that have co-evolved with their murine hosts ^7,12^. Previous studies have used re-wilded mouse models to study naturally occurring fungi and found that fungal colonization in re-wilded mice led to sustained neutrophil expansion and protected against bloodborne bacterial infection with *Staphylococcus aureus*^7^. In a comparative study of wildlings, wild mice, and conventional laboratory mice, wildlings and wild mice exhibited a much higher proportion of fungal DNA compared to the total DNA present than their conventional lab counterparts^8^. Furthermore, the composition of fungal communities differed: wildlings and wild mice had markedly more *Ascomycota* and much less *Basidiomycota* than conventional laboratory mice. In this study, we sought to isolate and characterize commensal fungi from pet store mice to elucidate their unique interactions with the host. Pet store mice carry a diverse and complex microbial community, including microbes and pathogens similar to those found in more natural, less sanitized environments. Laboratory mice cohoused with pet store mice become "dirty" mice, experiencing a natural-like microbial environment that results in mature and more human-like immune responses^9^. These features enable novel research opportunities to identify previously unrecognized host-microbe and microbe-microbe interactions. The multitude of microbiota transmitted—including viruses, bacteria, fungi, and eukaryotic organisms—facilitates the study of microbial succession, competition, synergy, and cross-kingdom interactions within the host, with observable impacts on immune cell phenotypes, cytokine production, and pathogen resistance. By scoping the gut microbiome of pet store mice from geographically distinct regions, we identified *Kazachstania pintolopesii* as the dominant fungal species. *Kazachastania* spp, were recently also characterized from laboratory as well as wild-caught mice^13,14^. Through a series of *in vivo* experiments and multi-omics analyses, we investigated the colonization dynamics and host immune responses upon the pet store *K. pintolopesii* strain colonization in conventional, antibiotics-treated, GF, and gnotobiotic laboratory mice. Like the previous studies, we found that *K. pintolopessii* strains isolated from pet store mice had superior ability to colonize laboratory mice irrespective of commensal bacteria rapidly outcompeting non-native fungal species. Additionally, we established that pet store-derived *K. pintolopesii* displays a benign immunological signature, distinct from *Candida albicans*, a human fungal commensal. This discovery marks the fungal commensalism, opening avenues for exploring how such interactions influence immune responses and microbial competition and sets a benchmark for commensal mycobiota.

## Results

### *K. pintolopesii* is the dominant gut fungus in pet store mice

SPF laboratory mice are maintained in controlled environments, shielded from potential pathogen exposure, and typically harbor microorganisms with limited diversity. In contrast, pet store mice live in large breeding colonies and are routinely exposed to rich varieties of microbes, including pathogenic bacteria, viruses, parasites, and fungi ^9^. As an example of this, electron microscopic imaging of the gut epithelial surface of laboratory mice revealed no significant epithelium-attached microbes. In contrast, the epithelial surfaces of pet store mice harbored many microbial-like structures of varying shapes and sizes (∼2-15 μm) (Fig. 1A). Therefore, we used pet store mice as a source to screen for commensal murine fungi.

**Fig. 1.**
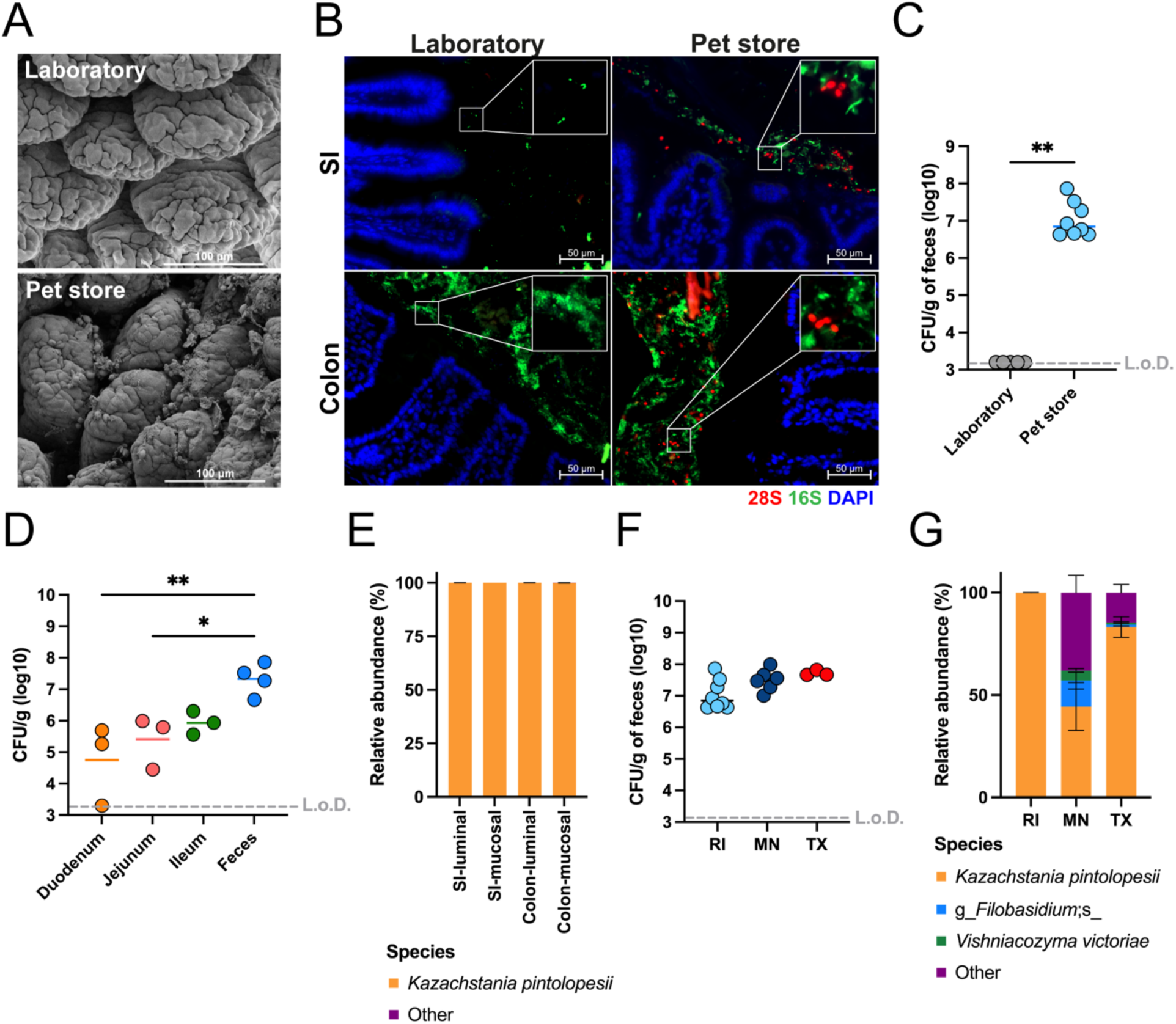
*K. pintolopesii* is the dominant fungus in the gut of pet store mice. (A) Representative scanning electron microscopy (SEM) image of the small intestine of laboratory mice and pet store mice. (B) Representative fluorescence *in situ* hybridization (FISH) images of fungal colonization in the small intestine and the colon of laboratory mice and pet store mice. Tissues were stained with DAPI (nuclei; blue), EUB338 probe (bacteria; green), and 28S probe (fungi; red). (C) Fungal burden in the feces of laboratory mice (*n* = 5) and pet store mice (*n* = 8). (D) Fungal burden in the intestinal tract of the pet store mice (*n* = 3 for tissues and *n* = 4 for feces). (E) Bar plot of relative abundance of fungal species from the ITS-marker gene sequencing at different sections of the intestine – luminal and mucosal part of the small intestine and the colon (*n* = 4 for SI-mucosal and *n* = 5 for other samples). Fungal burden (F) (RI, *n* = 8; MN, *n* = 6; TX, *n* =3) and bar plot of relative abundance of fungal species (G) in the feces of pet store mice (RI, *n* = 5; MN, *n* = 6; TX, *n* =8) in Rhode Island (RI), Minnesota (MI), and Texas (TX) in the USA. Bars indicate mean ± SEM. *p < 0.05 and **p < 0.01. L.o.D., limit of detection.

Using ribosomal RNA-based probes, we were able to find presence of fungal species among these epithelial-associated microbes in pet store mice, which are largely absent in laboratory mice (Fig.1B). Fungal burden in the feces of pet store mice was notably high (∼10^7^ CFU/g of feces), while no fungal populations were detectable in the feces of laboratory mice in our facility (Fig. 1C). Fungal colonization was observed throughout the gastrointestinal tract of the pet store mice, from the duodenum to the feces, with the fungal burden progressively increasing toward the caudal end of the intestine indicating that pet store mice are heavily colonized with fungi throughout the gastrointestinal tract (Fig. 1D).

To identify the commensal fungal species in the murine gut, we cultured the gut contents of the pet store mice for 5 days at 30 °C and randomly picked 26 colonies from the plates. Sequencing of the internal transcribed spacer (ITS) region of these colonies revealed that all colonies are the same fungal species, *Kazachstania pintolopesii* (Fig. S1A). The isolate appeared as a budding yeast and had an ellipsoidal shape with typical yeast cell structures such as a thick cell wall, nucleus, vacuole, and mitochondria (Fig. S1B and C). The isolate showed higher ITS sequence similarities with other *Kazachstania* species and *Saccharomyces cerevisiae*, whereas lower similarities with *Candida* and *Aspergillus* species (Fig. S1D). We also directly sequenced the ITS1 region of the gut metagenome of pet store mice. Intriguingly, this culture-independent analysis also showed that the intestines of pet store mice were predominantly colonized by a single fungal species, *K. pintolopesii*, which accounted for over 99% of the total sequence reads in both the luminal and mucosal regions of the small intestine and colon (Fig. 1E).

To determine whether this observation is specific to pet stores in a particular region (Rhode Island), fecal samples were collected from pet store mice in other geographical locations, including the Midwest United States (Minnesota) and the Southern United States (Texas) (Fig. S1E). Higher fungal burden was observed across all regions (Fig. 1F), with *K. pintolopesii* consistently identified as the predominant fungal species in the gut, regardless of the region (Fig. 1G). These findings indicate that *K. pintolopesii* is not limited to a specific geographical area but rather is a prevalent member of the mouse mycobiome under non-laboratory settings. It colonizes the gastrointestinal tract extensively, achieving high population levels both near the mucosal surface and within the intestinal lumen.

### *K. pintolopesii* stably colonizes laboratory mice irrespective of gut bacteria and provides colonization resistance against non-murine fungi

To test whether this fungal isolate from the pet store mice could colonize the intestines of the laboratory mice, we orally administered the isolated *K. pintolopesii* to laboratory mice. The isolated fungus colonized throughout the gastrointestinal tract, from the esophagus to the feces, of the laboratory mice (Fig. 2A). Furthermore, a high fungal burden (∼10^7^ CFU/g in the feces) was maintained for at least 6 weeks (Fig. S2A). The colonized *K. pintolopesii* was located closely to the gut epithelium of mice, co-localizing with bacteria, similar to the fungal colonization pattern observed in the pet store mice (Fig. S2B).

**Fig. 2.**
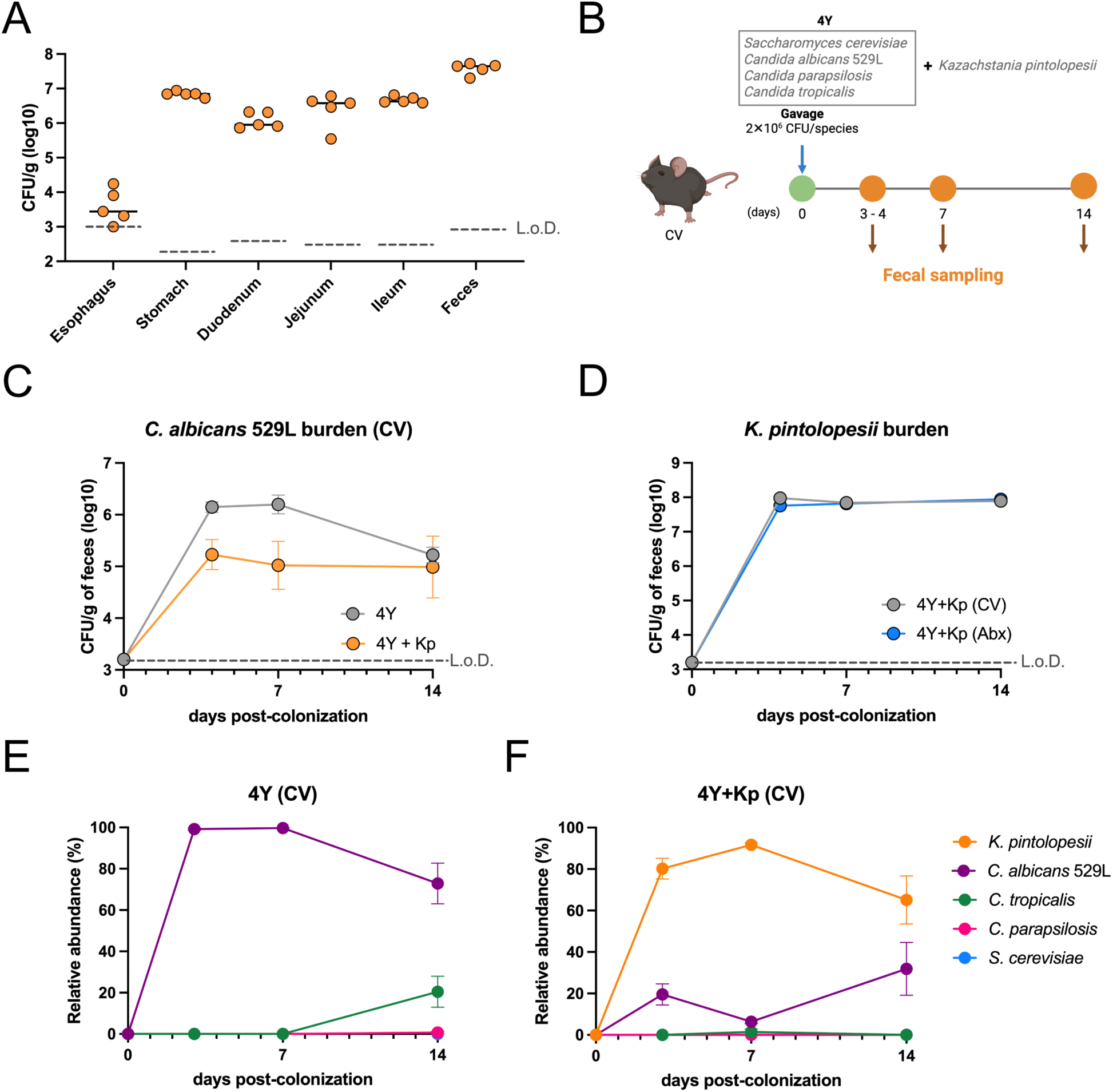
*K. pintolopesii* stably colonizes laboratory mice and provides colonization resistance against non-murine fungi. (A) Fungal burden in the gastrointestinal tract of mice at 14 days post-colonization of *K. pintolopesii* (*n* = 5). (B) Scheme of fungal competition assay in the conventional mice. (C) Burden of the *C. albicans* 529L in conventional mice with (4Y + Kp) or without *K. pintolopesii* (4Y). (D) Burden of the *K. pintolopesii* in conventional and antibiotic-treated mice. Line plot of relative abundance of fungal species from ITS-marker gene sequencing of the feces at different time points (days 3, 7, and 14) in conventional mice without (E) or with *K. pintolopesii* (F) (*n* = 5 per group). Dots indicate mean ± SEM. L.o.D., limit of detection. CV, conventional mice. Abx, antibiotic-treated mice. 4Y, four different yeast strains. Kp, *K. pintolopesii*-colonized mice.

The laboratory mice from various vendors have different bacterial communities in the gut ^15,16^. To test the colonization of *K. pintolopesii* in different bacterial compositions, we colonized the fungal isolate in mice bred at Brown University (Brown), Jackson Laboratory (Jax), and Taconic Biosciences (Tac). A high burden of *K. pintolopesii* (∼10^7^ CFU/g) was observed in the feces of Brown, Jax, and Tac mice (Fig. S2C). GF mice showed a comparable level of *K. pintolopesii* fecal colonization to other laboratory mice (Fig. S2D), demonstrating that the presence of commensal bacteria does not influence *K. pintolopesii*’s intestinal colonization. We performed 16S rRNA gene sequencing to assess the impact of *K. pintolopesii* colonization on the gut bacterial community, and *K. pintolopesii* colonization maintained bacterial community diversity, composition, and load without significant alterations (Fig. S2E-H). The ability of *K. pintolopesii* to colonize the gastrointestinal tract to high levels without impacting the bacterial community is in contrast to other fungal species that often require microbiome manipulation to establish colonization ^4,5,17–19^.

Recent studies showed that murine *Kazachstania* strains inhibited the other fungal growth in the mouse intestines ^13,14^. To examine whether the *K. pintolopesii* isolate affects the colonization of non-murine-specific fungi, a competition assay was conducted. The assay included five fungal species: *K. pintolopesii* and four other fungal species - *S. cerevisiae* BY4717*, C. albicans* SC5314*, C. parapsilosis* ATCC22019, and *C. tropicalis* MYA-3404 ^18–22^ (Fig. S3A). An equal number of viable fungal cells (∼2 × 10^6^ CFU/species) were orally administered simultaneously to both conventional mice and GF mice, and fecal samples were collected, followed by ITS sequencing. Notably, *K. pintolopesii* exhibited outstanding colonization ability, out-competing the other fungal species in both conventional and GF mice (Fig. S3B and C). In conventional mice, *K. pintolopesii* rapidly dominated the fungal community, accounting for over 95% of the total fungal population within 3 days of treatment (Fig. S3B). In GF mice, *C. albicans* SC5314 was initially the dominant species, with *C. tropicalis* MYA-3404 comprising over 10% of the population until day 7. However, the proportion of *K. pintolopesii* increased over time, eventually representing approximately 75% of the total fungal population by days 14 and 22 (Fig. S3C).

*C. albicans* SC5314 strain, the standard laboratory isolate, was used for the competition assay, but this strain is not murine-colonizing in the presence of commensal bacteria. In contrast, *C. albicans* 529L is a clinical isolate from the oral cavity and colonizes the murine gut without antibiotic treatment ^23,24^. We conducted another competition assay with *C. albicans* 529L to test whether *K. pintolopesii* can outcompete other fungal species in the presence of a murine-colonizing strain (Fig. 2B). In the absence of *K. pintolopesii*, *C. albicans* 529L was the most abundant fungal species and stably colonized in conventional mice (Fig. 2C and E). However, the addition of *K. pintolopesii* lowered the *C. albicans* 529L burden and outcompeted other fungal members (Fig. 2C, D, and F). In antibiotic-treated mice, the burden of *C. albicans* 529L was lower in the presence of *K. pintolopesii,* and the proportion of *K. pintolopesii* DNA was the most abundant among the fungal members at day 14 (Fig. S3D–G).

This suggests that *K. pintolopesii* is capable of stable colonization in laboratory mice, independent of the presence of commensal gut bacteria, and provides colonization resistance against non-murine fungi.

### *K. pintolopesii* colonization does not induce classical anti-fungal immune responses, including expansion of neutrophils and the Th17 population

Fungal colonization in the gut has been reported to trigger immune responses, including the expansion of neutrophils and T helper 17 cells (Th17) populations. *C. albicans*, a well-studied human commensal fungus, is known to induce neutrophils and Th17 populations in humans and mice ^25,26^. Rewilded mice, which harbor an increased fungal population in the gut, exhibited increased circulating neutrophil levels ^7,12^. In addition, mucosa-associated fungi induced type 17 immune responses ^27^. Interestingly, it was reported that intestinal colonization of *Kazachstania heterogenica* var. *weizmannii,* another murine-derived *Kazachstania* species, does not induce systemic neutrophilia and Th17 populations in lymph nodes, unlike *C. albicans* ^13^. To evaluate whether *K. pintolopesii* also induces the aforementioned classical anti-fungal immune responses, we measured the levels of neutrophils in blood upon the *K. pintolopesii* colonization. Interestingly, *K. pintolopesii* colonization did not induce circulating neutrophil levels in the peripheral blood (Fig. 3A). To mimic the systemic fungal dissemination, we introduced *K. pintolopesii* intravenously in mice, comparing it with *C. albicans* systemic infection (Fig. 3B). As expected, *C. albicans*-infected mice exhibited a poor survival rate and a high fungal burden in the kidneys, along with significant upregulation of *Il17a* gene expression in the kidney. Intriguingly, *K. pintolopesii*-infected mice did not show any pathological symptoms. All *K. pintolopesii*-infected mice survived and showed low to no detectable levels of fungal burden in the kidney. Additionally, *Il17a* gene expression in the kidney of *K. pintolopesii*-infected mice was significantly lower, approximately 14-fold lower than that observed in *C. albicans*-infected mice. (Fig. 3C – E). To assess neutrophils’ role in *K. pintolopesii* clearance during systemic infection, we depleted neutrophils using anti-Ly-6G (1A8 clone) antibody before *K. pintolopesii* infection (Fig. 3F). Despite complete neutrophil depletion, all mice survived without detectable kidney fungal burden (Fig. 3G-H and S4).

**Fig. 3.**
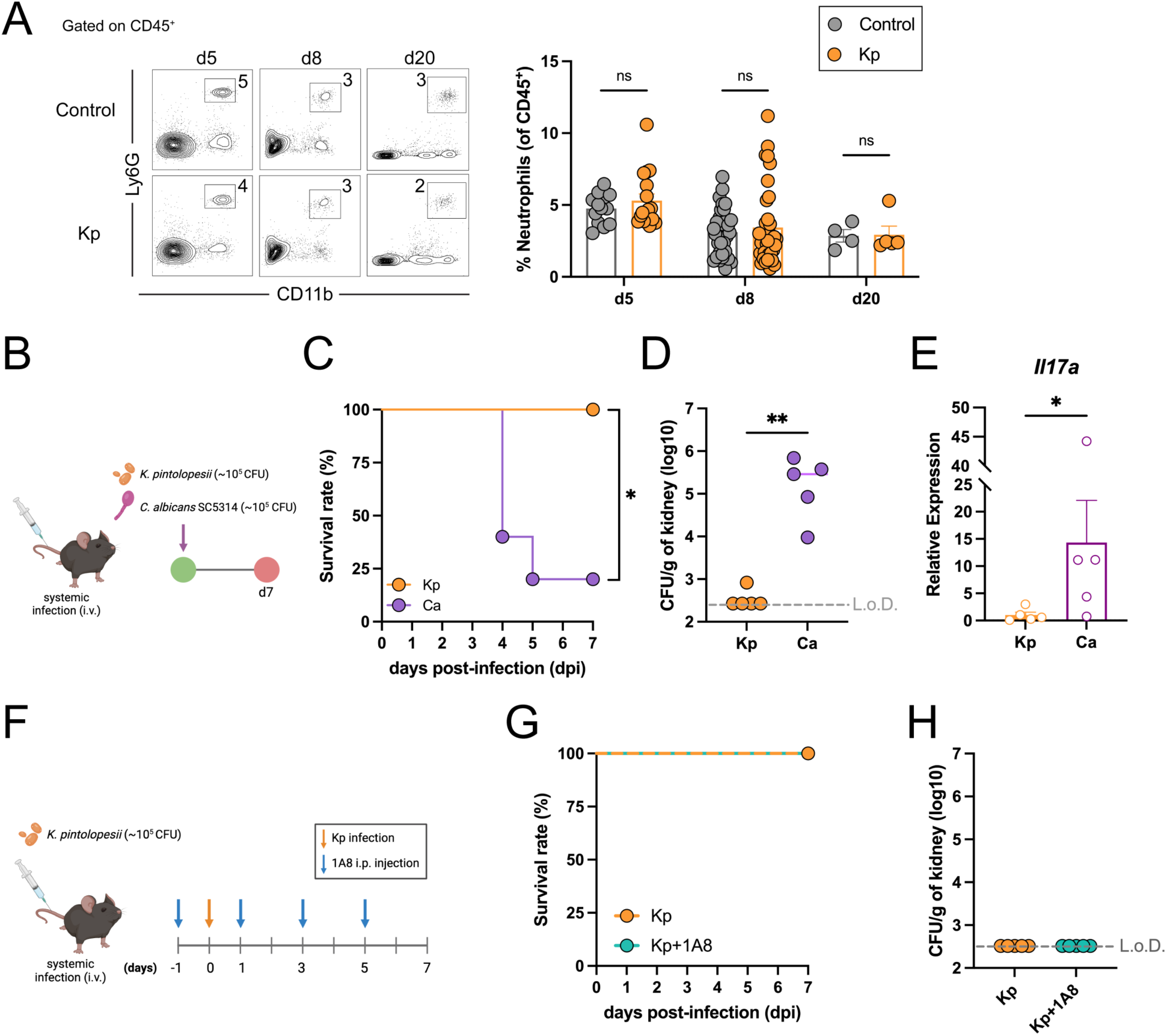
The systemic challenge with *K. pintolopesii* is efficiently resolved in a neutrophil-independent manner. (A) Representative flow cytometry plots and frequency of circulating neutrophils in the peripheral blood at 5 (Control, *n* = 13; Kp, *n* = 17), 8 (Control, *n* = 30; Kp, *n* = 35), and 20 days (Control, *n* = 4; Kp, *n* = 5) post-colonization of *K. pintolopesii*. Neutrophil populations were gated as CD45^+^ B220^-^CD11b^+^ Ly6G^+^. (B) Scheme of the systemic challenge with *K. pintolopesii* or *C. albicans* (*n* = 5 per group). (C) The survival curve after *K. pintolopesii* or *C. albicans* systemic infection. Fungal burden (D) and *Il17a* gene expression (E) in the kidney after *C. albicans* or *K. pintolopesii* infection. (F) Experimental schedule of neutrophil depletion and *K. pintolopesii* infection (*n* = 5 per group). (G) The survival curve after *K. pintolopesii* systemic infection with or without neutrophil depletion. (H) Fungal burden in the kidney after *K. pintolopesii* systemic infection with or without neutrophil depletion. Bars indicate mean ± SEM. *p < 0.05 and **p < 0.01. L.o.D., limit of detection. Kp, *K. pintolopesii*-colonized mice. Ca, *C. albicans*-colonized mice.

To assess the contribution of the adaptive immune system in colonization of commensal fungi, we introduced *K. pintolopesii* to Rag1 knock-out (KO) mice (Fig. 4A). Neither the body weight nor the fungal burden differed between wild-type (WT) and Rag1 KO mice (Fig. 4B and C). To assess the potential for systemic fungal dissemination and immune activation in the absence of B and T lymphocytes, we measured the number of neutrophils in the spleen. No significant differences were observed in neutrophil density between the two genotypes, indicating that the adaptive immune arms are neither engaged nor play a significant role in limiting *K*. *pintolopesii* burden (Fig. 4D). We also measured the levels of Th17, another key anti-fungal immune cell in mucosal immunity, following *K. pintolopesii* colonization. Interestingly, *K. pintolopesii* colonization did not increase Th17 cells in the small intestinal lamina propria (Fig. 4E).

**Fig. 4.**
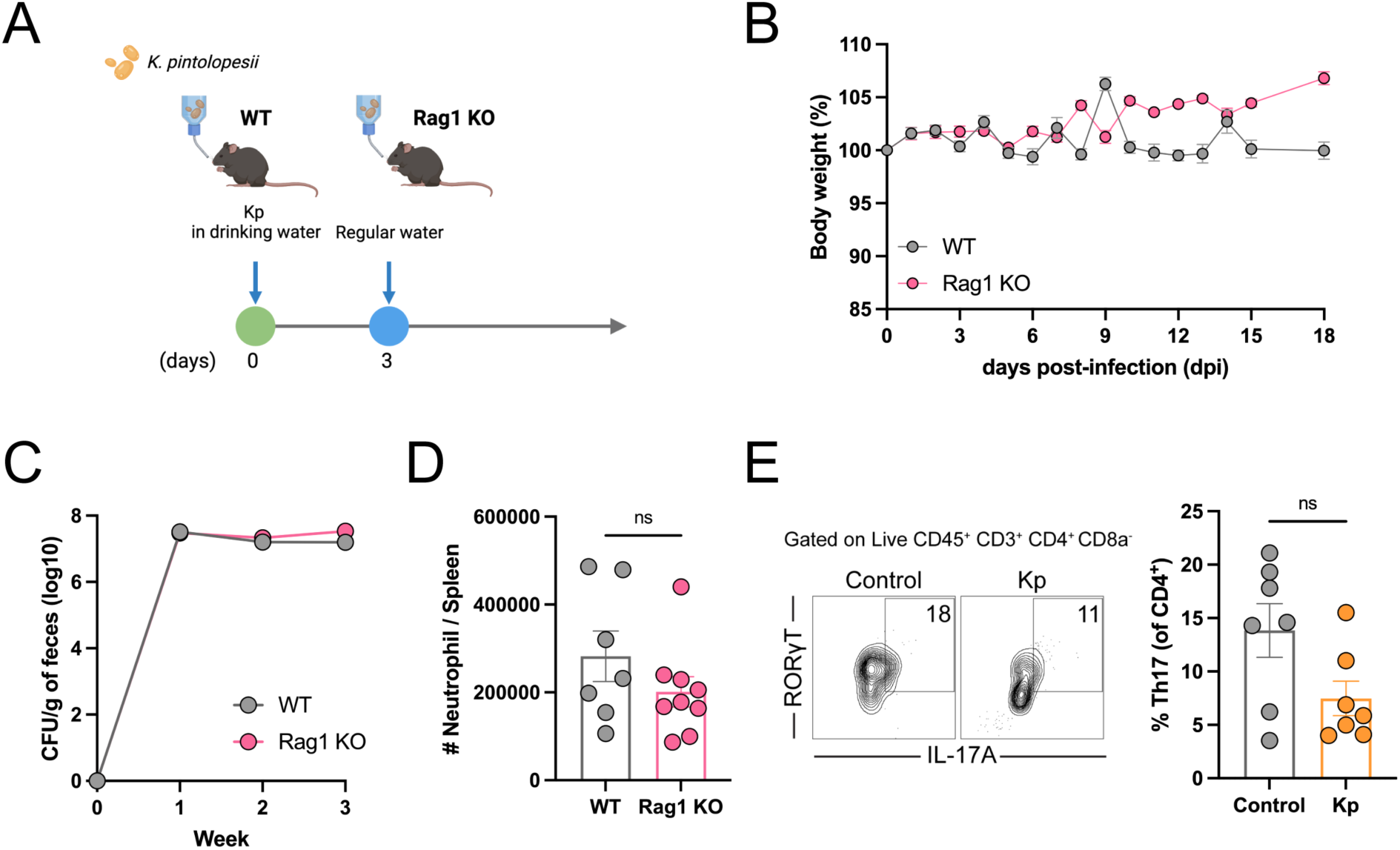
*K. pintolopesii* colonizes the intestine of mice in an adaptive immunity-independent manner. (A) Scheme of the *K. pintolopesii* colonization in wild-type (WT, *n* = 13) and adaptive immunity-deficient Rag1 knock-out mice (Rag1 KO, *n* = 18). (B) Body weight changes of WT and Rag1 KO mice after *K. pintolopesii* colonization. (C) Fungal burden in the feces of WT and Rag1 KO mice. (D) The number of neutrophils in the spleen at 19 days post-colonization of *K. pintolopesii* (WT, *n* = 7; Rag1 KO, *n* = 9). (E) Representative flow cytometry plots and frequency of Th17 cells in the small intestinal lamina propria at 14 days (*n* = 7 per group) post-colonization of *K. pintolopesii*. Th17 populations were gated as live CD45^+^ CD3^+^ CD4^+^ CD8a^-^ IL-17A^+^ RORγT^+^. Bars indicate mean ± SEM. Kp, *K. pintolopesii*-colonized mice.

The intestinal *C. albicans* colonization protects the host against systemic *C. albicans* infection ^26^. To assess whether the commensal fungus provides a similar cross-species protection to the host, *C. albicans* was systemically introduced into *K. pintolopesii*-colonized mice (Fig. 5A). However, no differences in survival rates or fungal burden in the kidneys were observed between the control and *K. pintolopesii*-colonized mice following systemic *C. albicans* infection (Fig. 5B and C). In contrast, we observed that the intestinal colonization by *C. albicans* in antibiotics-treated mice protected the host from systemic candidiasis (Fig. 5D – F).

**Fig. 5.**
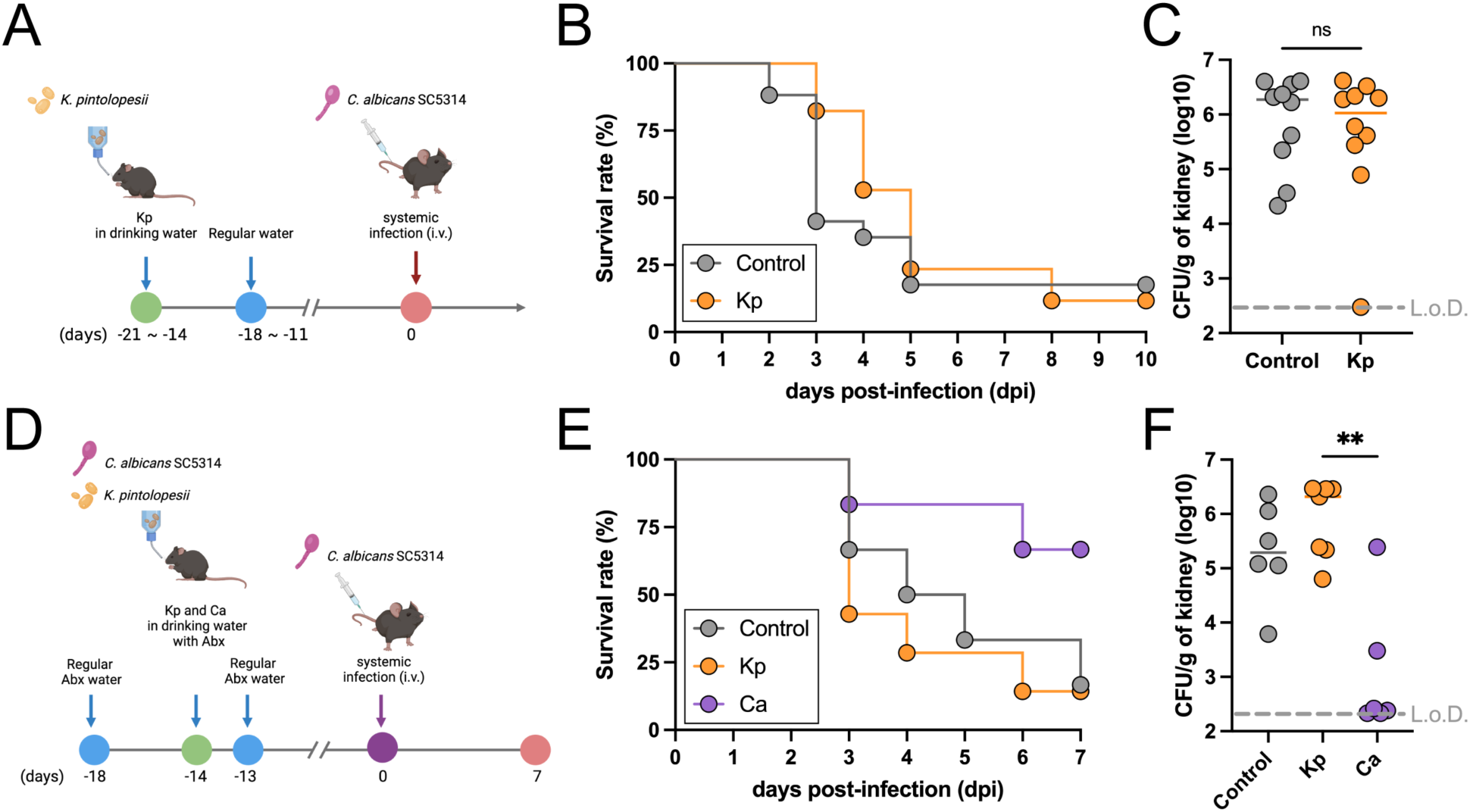
Intestinal colonization of *K. pintolopesii* does not protect the host against systemic *C. albicans* challenge. (A) Scheme of the *C. albicans* systemic challenge in the *K. pintolopesii*-colonized mice. Survival rate (B) (*n* = 17 per group) and fungal burden in the kidney (C) (*n* = 10 per group) after *C. albicans* challenge in control and *K. pintolopesii*-colonized mice. (D) Scheme of the *C. albicans* systemic challenge in the *K. pintolopesii*- or *C. albicans*-colonized mice with antibiotics (Abx) treatment. Survival rate (E) and fungal burden in the kidney (F) after *C. albicans* challenge in control (*n* = 6), *K. pintolopesii*-colonized (*n* = 7), and *C. albicans*-colonized mice (*n* = 6). Kp, *K. pintolopesii*-colonized mice. Ca, *C. albicans*-colonized mice.

These results indicate that *K. pintolopesii* does not trigger classical anti-fungal defenses, including neutrophilia and Th17 response, which have been previously reported in response to other *Kazachstania* strains ^13,14^. And the complete absence of the adaptive arm of the immune system does not enhance fungal pathogenicity. When *K. pintolopesii* was introduced systemically, the host was able to control the fungus via neutrophil-independent mechanisms, suggesting a unique host-fungal relationship that differs from typical anti-fungal immune responses.

### *K. pintolopesii* colonization induces type 2 immune responses in the gut and protects the host against roundworm infection

Although we examined the effects of *K. pintolopesii* colonization on the host immune system, we primarily focused on dominant anti-fungal immune responses, specifically neutrophils and Th17 cells. However, to thoroughly investigate the impact of *K. pintolopesii* colonization on the host’s intestine in an unbiased manner, we collected ileum samples 14 days post-colonization and performed bulk RNA sequencing. *K. pintolopesii* colonization showed significant transcriptional changes with 602 up-regulated genes and 1,213 down-regulated genes compared with control mice (Fig. S5). Notably, tuft cell and goblet cell marker genes were remarkably upregulated following *K. pintolopesii* colonization (Fig. 6A). To validate the RNA sequencing data, we quantified the number of tuft cells and goblet cells in the small intestine. As we saw in RNA sequencing data, the number of tuft cells and goblet cells was significantly higher in *K. pintolopesii-*colonized mice than in control mice (Fig. 6B and C). Tuft cells are a rare cell type in the intestinal epithelium, yet they play a pivotal role in goblet cell hyperplasia and in initiating type 2 immune responses ^28,29^. So, next examined *K. pintolopesii* also modulates other type 2-associated cells. We observed significant upregulation of T helper 2 (Th2) and group 2 innate lymphoid cells (ILC2s) by *K. pintolopesii* colonization (Fig. 6D and S6A).

**Fig. 6.**
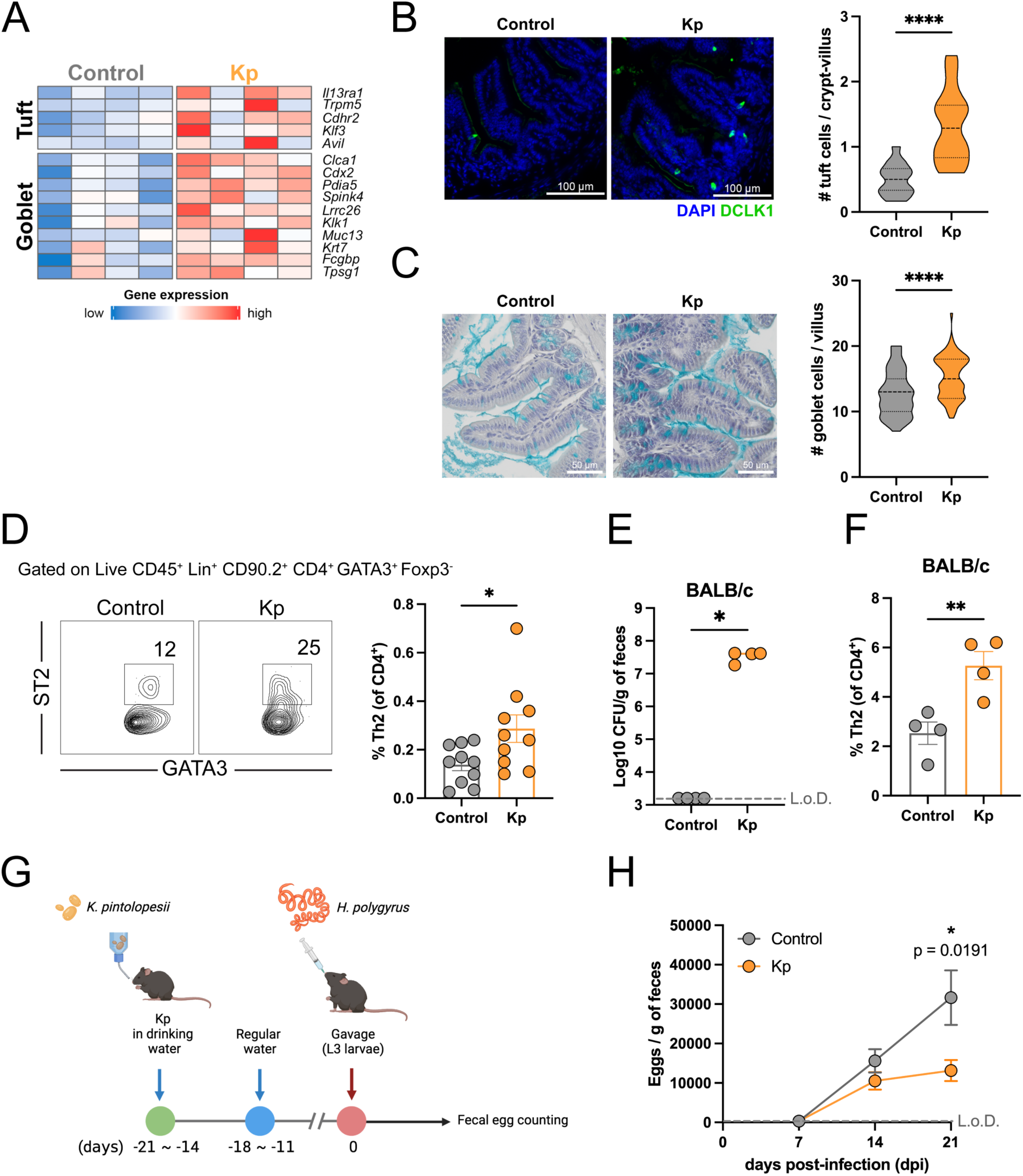
*K. pintolopesii* colonization increases the tuft cell population and induces type 2 immune responses in the gut and protects the host against *H. polygyrus* infection. (A) The heat map of expression of the tuft cell and the goblet cell marker genes in the small intestine (*n* = 4 per group). (B) Representative image and the number of tuft cells in the small intestine of control and *K. pintolopesii*-colonized mice. Tissues were stained with DAPI (nuclei; blue) and DCLK1 (tuft cells; green). 1-7 images were used for tuft cell counting per mouse, with 4–6 mice per group. (C) Representative image and the number of goblet cells in the small intestine of control and *K. pintolopesii*-colonized mice. Tissues were stained with hematoxylin (nuclei) and alcian blue (goblet cells). 6-15 villi were chosen for goblet cell counting per mouse, with 4–5 mice per group. (D) Representative flow cytometry plots and frequency of Th2 in the small intestinal lamina propria of control and *K. pintolopesii*-colonized mice (*n* = 10 per group). Th2 populations were gated as live CD45^+^ Lin^+^ CD90.2^+^ CD4^+^ GATA3^+^ Foxp3^-^ ST2^+^. Lin: CD3ε, CD11b, Ly-6G/Ly-6C, CD19. (E) Fungal burden in the feces of BALB/c mice after *K. pintolopesii* colonization (*n* = 4 per group). (F) Frequency of Th2 in the small intestinal lamina propria of control and *K. pintolopesii*-colonized BALB/c mice (*n* = 4 per group). (G) Scheme of the *H. polygyrus* challenge. (H) The quantification of worm eggs in the feces of the control and *K. pintolopesii*-colonized mice (Control, *n* = 9; Kp, *n* = 10). Bars indicate mean ± SEM. *p < 0.05, **p < 0.01, and ****p < 0.0001. Kp, *K. pintolopesii*-colonized mice.

To check if this induction of type-2 immunity is not limited only to the C57BL/6 mouse strain, we colonized *K. pintolopesii* in the prototypical Th2-type mice, BALB/c ^30,31^. Colonized BALB/c mice exhibited a high fungal burden (Fig. 6E) and significantly increased frequencies of Th2 and ILC2 levels in the small intestine, similar to what was observed in C57BL/6 mice (Fig. 6F and S6B). However, circulating neutrophil levels and the Th17 population in the small intestine were comparable to those in control mice (Fig. S6C and D).

Next, to assess the functional relevance of this type-2 immunity, we challenged control and *K. pintolopesii*-colonized mice with a murine intestinal nematode, *Heligmosomoides polygyrus*, that is controlled by type-2 immune responses (Fig. 6G). The number of eggs in the fecal samples was significantly lower in *K. pintolopesii*-colonized mice compared with control mice at 21 days post-infection (Fig. 6H). This suggests that *K. pintolopesii* colonization in the mouse intestine protects the host from a parasitic nematode.

## Discussion

Naturalized, wilded, and pet store-cohoused mice have emerged as valuable models for studying the impact of microbes encountered outside the laboratory setting. These mice provide researchers with a more accurate representation of the complex microbial interactions that occur in natural environments, offering insights that conventional laboratory mice cannot. Transfer of gut microbiome from wild mice to the GF mice exhibited improved protection against viral infection, and embryo transfer of laboratory mice into wild mice showed human-like immune phenotypes ^8,32^. Exposing laboratory mice to wild environments and then recapturing them is another way to colonize the wild microbiome in laboratory mice. These rewilded mice exhibit increased circulating neutrophils and enhanced granulopoiesis ^7,12^. Pet store mice are another valuable resource for studying host-microbe interactions in mice. Although they do not live in the natural environment like wild mice, they are exposed to the dirty environment in which humans live. When laboratory mice are exposed to the pet store mice microbiome by co-housing, co-housed laboratory mice harbor various microbes, such as viruses, fungi, and worms, and exhibit a similar immune composition to that of human adults ^9,33^.

While studies have explored interactions between mice and their complex microbiomes in toto, the specific relationships between hosts and individual microbial kingdoms are less explored, especially in the presence of diverse pathogenic groups. Among various microbes, we focused on the role of commensal fungi in the murine gut. Fungi ubiquitously exist in various environments, including multiple body sites of mammals ^34^. In general, the importance of commensal fungi is underestimated compared to commensal bacteria due to fewer available databases and lower cell numbers compared to bacteria in the gut. However, fungi can contribute unique metabolic functions as eukaryotes, and even a minor commensal member can affect host phenotypes in the gut ^15,35,36^. In our study, we aimed to identify murine commensal fungi that will allow us to take advantage of the robust mouse model to study fungal commensalism. We successfully isolated *K. pintolopesii* from the intestines of pet store mice, which constituted the dominant fungal species in the pet store mice regardless of geographic location. In laboratory settings, *K. pintolopesii* was capable of stable colonization without any manipulation of the gut microbiome and was not harmful to the host.

Two recent studies also identified *Kazachstania* species from immunodeficient and wild mice that can colonize the gut of laboratory mice to high numbers. While the study by Kralova et al. investigates *Kazachstania heterogenica* var. *weizmannii* isolated from an immunodeficient mouse model ^13^, the study by Liao et al. examines *K. pintolopesii* isolated from free-living mice ^14^. The *Kazachastania* sp. in all three studies demonstrates a remarkable ability to colonize the murine gastrointestinal tract and outcompete or prevent colonization by non-murine fungi, including *C. albicans*, showcasing their potential as powerful modulators of the gut mycobiome. Other studies using the wildling mice also support that *K. pintolopesii* is a dominant commensal fungus of the mouse intestine ^37,38^.

*K. weizmannii* exhibits a stealth colonization strategy, inducing minimal immune responses under normal conditions but mitigating *C. albicans*-induced pathology ^13^. In contrast, *K. pintolopesii* isolated in Liao et al. and our study triggers a robust type 2 immune response, enhancing resistance to helminth infections while exacerbating gastrointestinal allergy ^14^.

The Liao et al. study shows that *K. pintolopesii* triggers a type 2 immune response, characterized by eosinophilia and Th2 cell polarization, but only when the mucosal barrier is impaired (e.g., during fiber-free diet, antibiotic treatment, or in GF mice) ^14^. In contrast, we observed that the *K. pintolopesii* strain from pet store mice increases type 2 immune cells in the small intestine and provides cross-kingdom protection against helminth infection even without any need for barrier impairment, suggesting strain-level differences in *K. pintolopesii* mice strains between the two studies. Perhaps the microbial environment of pet store mice, which have a higher burden of parasitic worms and pathogens, has forced *K. pintolopesii* to induce type 2 immunity ^39–41^. Cross-kingdom interactions highlight the complexity and interconnectedness of microbial communities and their host organisms. These interactions can have significant implications for understanding disease resistance, developing new therapeutic strategies, and managing ecosystems.

In conclusion, our study underscores the critical importance of strain-specific, commensal fungal interactions in shaping the gut mycobiome and influencing host physiology. Future strain-level studies of commensal fungi will provide deeper insights into host-fungal commensalism. The discovery of the current study will lead to a more nuanced understanding of how commensal fungi contribute to host health, influence immune system development, and potentially protect against pathogens or modulate disease outcomes. This approach could reveal new therapeutic targets or probiotic strategies for improving gut health and managing various gastrointestinal disorders.

## Methods

### Mice

Conventional wild-type C57BL/6 mice were purchased from Jackson Laboratory and Taconic Biosciences or were bred in the animal facility at Brown University. BALB/c mice were purchased from Jackson Laboratory. Rag1 KO mice were used as an adaptive immunity-deficient mouse model. GF mice were wild-type C57BL/6 background, and they were raised and bred in flexible film isolators in a gnotobiotic facility at Brown University. Pet store mice were purchased from local pet stores. Fecal pellets of pet store mice in Minnesota and Texas were freshly delivered to Brown University and used for further experiments. Experiments were performed according to the protocols approved by the Institutional Animal Care and Use Committees of Brown University.

### Fungal isolation from the gut of pet store mice

Cecal contents and feces, collected from pet store mice, were suspended in sterile PBS and plated on Sabouraud Dextrose (SD) agar with streptomycin (40 mg/L), followed by incubation at 30 °C for 5 days. A total of 26 single colonies were randomly picked from the plates, and the ITS region of fungal colonies was amplified by colony PCR using the Phire Green Hot Start II PCR Master Mix (Thermo Scientific) with ITS1-F/ITS4 primer pair (ITS1-F: 5’-CTTGGTCATTTAGAGGAAGTAA-3’, ITS4: 5’-TCCTCCGCTTATTGATATGC-3’) ^42,43^. The amplification program consisted of 94 °C for 10 min, followed by 35 cycles of 94 °C for 30 sec, 52 °C for 30 sec, and 72 °C for 1 min, and 72 °C for 8 min. The amplicon was sequenced by Sanger sequencing (GENEWIZ), and MOLE-BLAST (Basic Local Alignment Search Tool) was performed against the ITS database of NCBI to identify the isolated colonies. The exported phylogenetic tree from MOLE-BLAST was visualized using the R package *ggtree*.

### Fungal culture

All fungal strains were cultured in SD or yeast extract-peptone-dextrose (YPD) media. The isolate, *K. pintolopesii*, was cultured at 37 °C for 1 day with shaking. This culture was inoculated in fresh media and cultured for 1 day under the same conditions. *S. cerevisiae* BY4717 was cultured at 30 °C for 2 days with shaking. *C. albicans* SC5314, *C. albicans* 529L, *C. parapsilosis* ATCC22019, and *C. tropicalis* MYA-3404 were cultured overnight at 30 °C with shaking. For systemic infection, the *C. albicans* overnight culture was inoculated in fresh media and cultured for 4 h under the same conditions.

### Fungal infection

For intestinal colonization of non-colonizing fungi, antibiotic water was prepared with penicillin (750 U/mL), streptomycin (1 mg/mL), and 2.5% (w/v) glucose. *C. albicans* cultures (∼10^6^ CFU/mL) were resuspended in antibiotic water and administered. The same antibiotic water was provided to mice in both the control and *K. pintolopesii* groups. All antibiotic waters were administered starting four days prior to oral inoculation. One day later, the fungal antibiotic water was replaced with fresh antibiotic water, and fresh antibiotic water was provided every three or four days throughout the whole experimental period. For systemic infections, *K. pintolopesii* (∼10^5^ CFU/200 μL) or *C. albicans* (∼10^5^ CFU/200 μL) was injected into mice via the retro-orbital venous sinus or lateral tail vein.

All fungal cultures were washed twice with sterile PBS and resuspended in the proper solution for infection.

The fecal pellets and the contents of the intestine were resuspended in sterile PBS with antibiotics (500 mg/L ampicillin, 250 mg/L streptomycin, 225 mg/L kanamycin) to determine fungal burden. The other tissues were homogenized in sterile PBS with antibiotics. All samples were serially diluted in sterile PBS with antibiotics and plated on SD agar with streptomycin (40 mg/L), followed by incubation at 30 °C or 37 °C for 1 or 2 days based on the optimal condition of each fungus.

### Fungal competition assay

For oral administration of various fungal species, each fungal culture was mixed with the same CFU and gavaged to conventional, antibiotics-treated mice, or GF mice (∼2 × 10^6^ CFU/species in 200 μL per mouse).

N-acetyl transferase (NAT)-integrated *C. albicans* 529L strain was used for the treatment. To determine *C. albicans* 529L burden, fecal samples were plated on YPD agar plates with streptomycin (40 mg/L) and nourseothricin sulfate (200 mg/L), followed by incubation at 30 °C for 1 day. Fecal *K. pintolopesii* burden was determined by the colony morphology on SD agar plates with streptomycin (40 mg/L), followed by incubation at 37 °C for 2 days.

### Immune cell isolation and flow cytometry

To isolate peripheral blood lymphocytes, blood samples were collected from the submandibular vein by cheek punch in heparin-containing polypropylene tubes. Red blood cells were removed from blood samples by two rounds of ACK (Ammonium-Chloride-Potassium) lysis buffer treatment. Lymphocytes from the spleen and the small intestinal lamina propria were isolated as described in Beura et al. ^44^ and Grizotte-Lake et al. ^45^.

Isolated cells were stained with the corresponding cocktail of fluorescently labeled antibodies (CD3, CD4, CD8a, CD11b, CD45, CD90.2, B220, Ly6C, Ly6G, ST2) and biotin-conjugated anti-mouse lineage (CD3ε, CD11b, CD19, Ly6G/Ly6C) and streptavidin. Cell viability was determined using Ghost Dye 780 or Fixable Viability Dye eFluor™ 450. After surface staining, cells were fixed and permeabilized, followed by intracellular staining with the corresponding cocktail of fluorescently labeled antibodies (Foxp3, GATA3, RORγT, IL-17A). The stained samples were acquired using the Aurora flow cytometer (Cytek) and were analyzed with FlowJo 10 software (BD).

### Neutrophil depletion

For neutrophil depletion, anti-Ly-6G antibody (1A8 clone; Biolegend) or isotype control (Biolegend) was administered intraperitoneally to mice every other day (200 μg in 200 μL per mouse) from the day before the *K. pintolopesii* challenge.

### DNA extraction for microbiome and mycobiome analysis

The small intestine (ileum) and colon were flushed with sterile PBS to collect luminal contents, and intestinal material was obtained by centrifugation at 5,000 rpm for 10 min. After the flush, the small intestine and colon were scraped and collected in sterile PBS, followed by centrifugation at 5,000 rpm for 10 min to get mucosal contents. Fecal samples were directly collected from the rectum. Samples (∼50 mg) were suspended in 50 mM Tris buffer (pH 7.5) supplemented with 1 mM EDTA, 0.2% β-mercaptoethanol, and 1000 units/mL of lyticase (Sigma-Aldrich), followed by incubation at 37 °C for 30 min. After incubation, genomic DNA was extracted from samples using a Quick-DNA Fecal/Soil Microbe Microprep Kit (Zymo Research) according to the manufacturer’s protocol and was stored at −20°C until further processing.

### Bacterial load measurement

Bacterial load in the feces was measured by quantitative real-time PCR (qPCR). Reactions were prepared using Maxima SYBR Green/ROX qPCR Master Mix (Thermo Scientific). Bacterial DNA contents in the intestine were determined by 16S rRNA gene contents using the 340F/514R primer pair (340F: 5’-ACTCCTACGGGAGGCAGCAGT-3’, 514R: 5’-ATTACCGCGGCTGCTGGC-3’). Bacterial load was expressed as a 2^-Ct^ value and normalized to the weight of the starting fecal material.

### 16S rRNA gene sequencing and bacterial community analysis

The V4/V5 region of the bacterial 16S rRNA gene was amplified using the Phusion High-Fidelity DNA polymerase (Thermo Scientific) with 518F/926R primer pair (518F: 5’-CCAGCAGCYGCGGTAAN -3’, 926R: 5’-CCGTCAATTCNTTTRAGT -3’), which were conjugated with overhang adapter sequences for Illumina MiSeq. The amplification program consisted of 95 °C for 3 min, followed by 35 cycles of 95 °C for 30 sec, 58 °C for 30 sec, and 72 °C for 30 sec, and 72 °C for 10 min. DNA libraries were constructed and sequenced on Illumina MiSeq 2 × 300 bp paired-end sequencing (Rhode Island Genomics and Sequencing Center).

The raw sequencing reads were processed using the QIIME2 pipeline ^46^. DADA2 ^47^ was used to perform denoising and to construct the amplicon sequence variants (ASVs) table. The pre-trained Naïve Bayes classifier on the SILVA 132 database ^48^ was used for taxonomic assignment. Singletons and all features annotated as mitochondria or chloroplast were removed from the table, and the abundance of bacterial taxa was expressed as a percentage of the total 16S rRNA gene sequences. For alpha and beta diversity, the feature table was rarefied to 10,434 reads. Shannon was used as an alpha diversity index, and principal coordinates analysis (PCoA) based on weighted UniFrac distances was used for beta diversity. The R package *qiime2R* (https://github.com/jbisanz/qiime2R) was used for visualization.

### ITS sequencing and fungal community analysis

The ITS1 region of the fungal DNA was amplified using the Phusion High-Fidelity DNA polymerase (Thermo Scientific) with ITS1-F/ITS2 primer pair (ITS1-F: 5’-CTTGGTCATTTAGAGGAAGTAA-3’, ITS2: 5’-GCTGCGTTCTTCATCGATGC -3’)^42,43^, which were conjugated with overhang adapter sequences for Illumina MiSeq. DNA library construction and sequencing were conducted following the same procedures for 16S rRNA gene sequencing.

The raw sequencing reads were processed using the QIIME2 pipeline ^46^. The reads were trimmed using Q2-ITSxpress ^49^. Trimmed sequence reads were denoised, and sequence variants were identified by DADA2 ^47^. Taxonomy was assigned using a pre-trained Naïve Bayes classifier on the UNITE v8.2 database ^50^. Singletons were removed from the table, and the abundance of fungal taxa was expressed as a percentage of total sequence reads.

### RNA extraction from tissues

Small pieces of the small intestine (ileum) and the kidney were stored in RNAlater (Invitrogen). Total RNA from tissues was extracted using PureLink™ RNA Mini Kit (Invitrogen) or RNeasy Plus Mini Kit (Qiagen) according to the manufacturer’s protocol and was stored at −20°C until further processing.

### Quantitative PCR (qPCR)

cDNA was synthesized from isolated RNA with M-MLV Reverse Transcriptase (Invitrogen). qPCR reactions were prepared using Maxima SYBR Green/ROX qPCR Master Mix (Thermo Scientific) with primer set for *Il17a* (F: ATCAGGACGCGCAAACATGA, R: TTGGACACGCTGAGCTTTGA) gene. Gene expression was normalized to *Gapdh* (F: TGGCAAAGTGGAGATTGTTGCC, R: AAGATGGTGATGGGCTTCCCG) and relative expression was calculated using the ^ΔΔ^Ct method.

### RNA-Seq

Extracted RNA samples were submitted for library construction and sequencing (Novogene). The constructed libraries were sequenced on the Illumina NovaSeq X Plus platform (Paired-end 150 bp). Adapter and low-quality reads were trimmed from the raw sequence reads using Trimmomatic ^51^. Trimmed sequence reads were aligned to the mm10 mouse genome using the STAR aligner ^52^. DESeq2 ^53^ was used to identify differentially expressed genes (DEGs) between groups when *P* < 0.05. The heatmap and volcano plot were drawn using the R packages Complex heatmap ^54^ and EnhancedVolcano (https://github.com/kevinblighe/EnhancedVolcano), respectively. The cell-specific gene list was obtained from PanglaoDB ^55^ and utilized as a cell marker gene set for gene set enrichment analysis (GSEA). The R package clusterProfiler ^56^ was used to perform GSEA.

### Fluorescence *in situ* hybridization (FISH)

The small intestine and the colon were fixed in Methacarn (Bioworld) and embedded in paraffin. Tissues were sectioned at 7 μm and deparaffinized in xylene followed by rehydration in 95% EtOH, 90% EtOH, and H_2_O. After deparaffinization and rehydration, tissue sections were hybridized with universal bacterial probes (Alexa 488-GCTGCCTCCCGTAGGAGT; Alexa 488-GCAGCCACCCGTAGGTGT) and a universal fungal probe (Alexa 555-CTCTGGCTTCACCCTATTC) directed against the 16S rRNA gene and 28S rRNA gene, respectively. DAPI was used to stain the nucleus.

### Intestinal cell staining

The small intestine was fixed in Methacarn (Bioworld) and embedded in paraffin. Fixed tissues were sectioned at 7 μm and deparaffinized in xylene followed by rehydration in 100% EtOH, 95% EtOH, 70% EtOH, and H_2_O.

For tuft cells, slides were incubated in citrate buffer (10 mM Trisodium citrate, 0.05% Tween 20, pH 6.0) for 20 min at 95 °C for antigen retrieval. Slides were blocked with 1% bovine serum albumin (BSA) followed by overnight incubation with rabbit anti-mouse DCAMKL1 (Abcam) antibodies at 4 °C. After incubation, slides were washed and incubated with goat anti-rabbit Alexa Fluor 546 (Invitrogen) secondary antibodies for 1 h at room temperature. Slides were counterstained with DAPI.

For goblet cells, slides were incubated with Alcian blue (StatLab) for 30 min, followed by washing in running tap water for 2 min. Slides were counterstained with hematoxylin for 2 min, followed by washing in running tap water for 30 sec. The slides were 10 dips in the clarifier and bluing reagent with washing in tap water, respectively. Slides were dehydrated in 95% EtOH, 100% EtOH, and xylene.

### Scanning electron microscopy (SEM) & transmission electron microscopy (TEM)

For SEM, *K. pintolopesii* cultures were washed twice with sterile PBS, and a thin layer was applied to the coverslips. The coverslips were fixed in fixation buffer (2.5% glutaraldehyde, 0.15 M sodium cacodylate, 2% paraformaldehyde, 2 mM calcium chloride, 0.1 M sucrose, pH 7.4), followed by incubation for 2 h at room temperature. For the small intestine, sections of the tissue were flushed with sterile PBS and then cut into 1 cm sections. They were incubated in the same fixation buffer as previously described. After incubation, the coverslips were washed three times with washing buffer (2 mM calcium chloride, 0.15 M sodium cacodylate, 0.1 M sucrose). After washing, samples were incubated overnight in 2% osmium tetroxide, 2.5% glutaraldehyde, 0.15 M sodium cacodylate, and 2% paraformaldehyde. Samples were washed with DI water three times and dehydrated in an EtOH gradient as follows: 20%, 50%, 70%, 90%, 95% and then 100%. Each EtOH step was incubated for 20 min at room temperature and then washed twice with 100% EtOH. Samples were flash dehydrated using a critical point dryer and then mounted on coverslips for imaging. Thermo Apreo SEM was used for imaging.

For TEM, *K. pintolopesii* cultures were pelleted in the 1.5 mL Eppendorf tubes. The fixation buffer was added to the tubes and incubated for 2 h at room temperature. The samples were then washed three times with the washing buffer. After washing, samples were incubated overnight in 2% osmium tetroxide, 2.5% glutaraldehyde, 0.15 M sodium cacodylate, and 2% paraformaldehyde. Samples were washed with DI water three times, followed by incubation overnight at 4 °C in saturated uranyl acetate solution. The next day, samples were dehydrated in the EtOH gradient as we described in SEM preparation and washed three times in 100% EtOH. Then, the samples were infiltrated with Hard Spurr’s resin and rocked on a rotator at room temperature. The gradient started at 25% Hard Spurr’s and 75% EtOH and gradually increased until 100% resin. Each step lasted 2 h, and the final 100% resin step was left overnight. The samples in resin were placed in resin molds and then let to harden for 18-24 h at 60 °C. The pellets in resin were cut with a knife into 100-200 nm sections and observed on PHILIPS 410 TEM.

### Heligmosomoides polygyrus infection

*H. polygyrus* L3 larvae were orally gavaged to mice (200 L3 larvae per mouse). After infection, fecal samples were collected every week. Fecal samples were resuspended in water and mixed with the same volume of supersaturated salt solution. The resuspended fecal solution was immediately loaded on a McMaster counting chamber (Eggzamin) and eggs were counted under the microscope. The counted eggs were normalized by fecal weight.

### Statistical analysis

All statistical analysis and plotting were performed on R (https://www.R-project.org) and Prism (GraphPad). The normality of data was assessed by the Shapiro-Wilk test. If data followed normal distribution, parametric tests (unpaired two-tailed t-test for two groups, One-way ANOVA with Tukey’s multiple comparisons test for more than two groups) were performed. If data did not follow normal distribution, non-parametric tests (Mann-Whitney test for two groups, Kruskal-Wallis test with Dunn’s multiple comparisons test for more than two groups) were performed to find significant differences between groups. For the survival curve, the log-rank (Mantel-Cox) test was used to find significant differences between groups. A two-way ANOVA test was used to find significant differences in body weight changes at each time point between the two groups. *p < 0.05, **p < 0.01, ***p < 0.001, ****p<0.0001.

## Data availability

Raw sequence reads from 16S and ITS sequencing have been deposited in the NCBI Sequence Read Archive (SRA) under accession number PRJNA1183864. Bulk RNA-seq data have been deposited in the NCBI GEO database under accession number GSE281418. This paper does not report the original code.

## Acknowledgments

Schematics were created with BioRender.com. We thank Dr. Richard Bennett (Brown University) for providing *S. cerevisiae* BY4717, *C. albicans* 529L, *C. albicans* SC5314, *C. parapsilosis* ATCC22019, and *C. tropicalis* MYA-3404 strains. GH was supported by an American Association of Immunologists Careers in Immunology Fellowship. NNJ is supported by the Damon Runyon Cancer Research Foundation (DRG-2427-21) and NIAID 1K22AI177360-01. SV is supported by R01DK113265, P20GM109035 (pilot project) and Brown University Seed grant. LKB is supported by Searle Scholar Foundation and this work is also supported by Brown University Seed grant. The Thermo Apreo VS SEM was purchased with a high-end instrumentation grant from the Office of the Director at the National Institutes of Health (S10OD023461).

## Author contributions

Conceptualization: SV, LKB, GH; Methodology: SV, LKB, GH, RY, MHH, NNJ; Investigation: SV, LKB, GH, RY, MHH, AB, KB, JP; Resources: SV, LKB, NNJ; Funding acquisition: SV, LKB; Supervision: SV, LKB; Writing – original draft: SV, LKB, GH; Writing – review & editing: SV, LKB, GH

