## Supplementary information for "A host-adapted commensal fungus from pet store mice drives type 2 immunity and cross-kingdom protection"

A

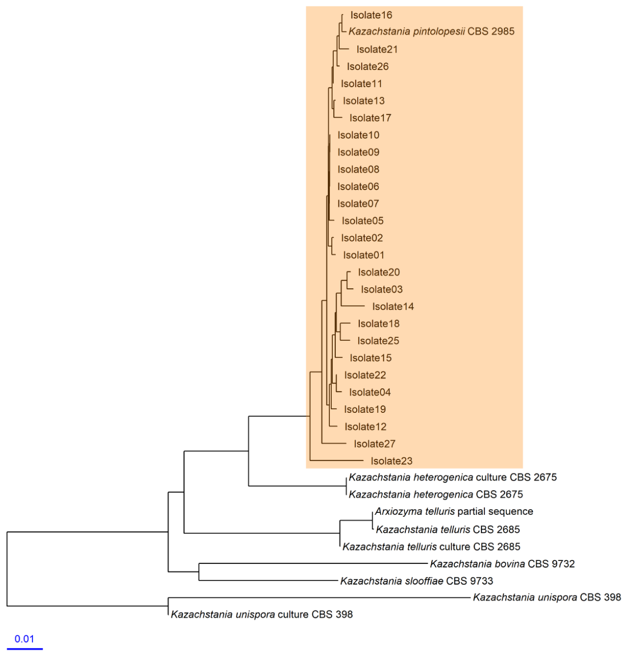


B

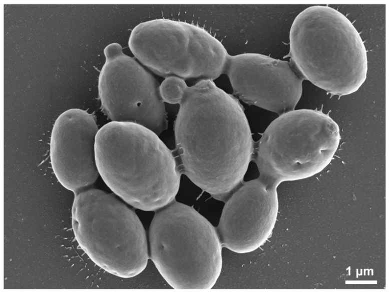


C

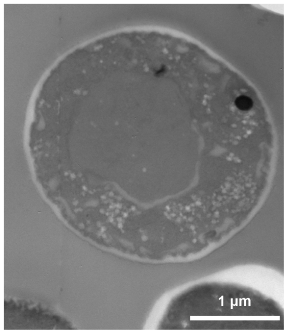


D

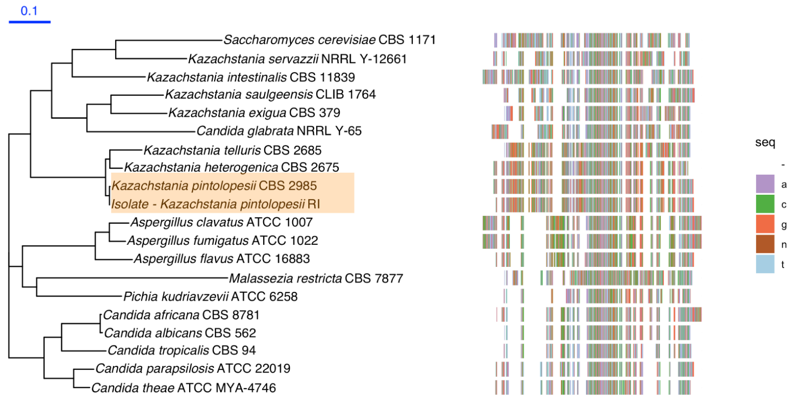


E

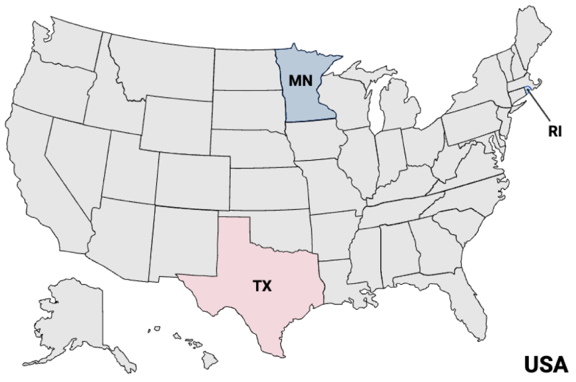

**Fig. S1. Identification of the isolated fungal strain from the pet store mice intestine.** (A) MOLE-BLAST result based on ITS sequences of isolates from the pet store mice intestine. Scanning electron microscopy (SEM) (B) and transmission electron microscopy (TEM) image (C) of fungal isolate. (D) Phylogenetic analysis of the isolate with various fungal species. (E) The United States map is used to designate the locations of fecal samples collected.

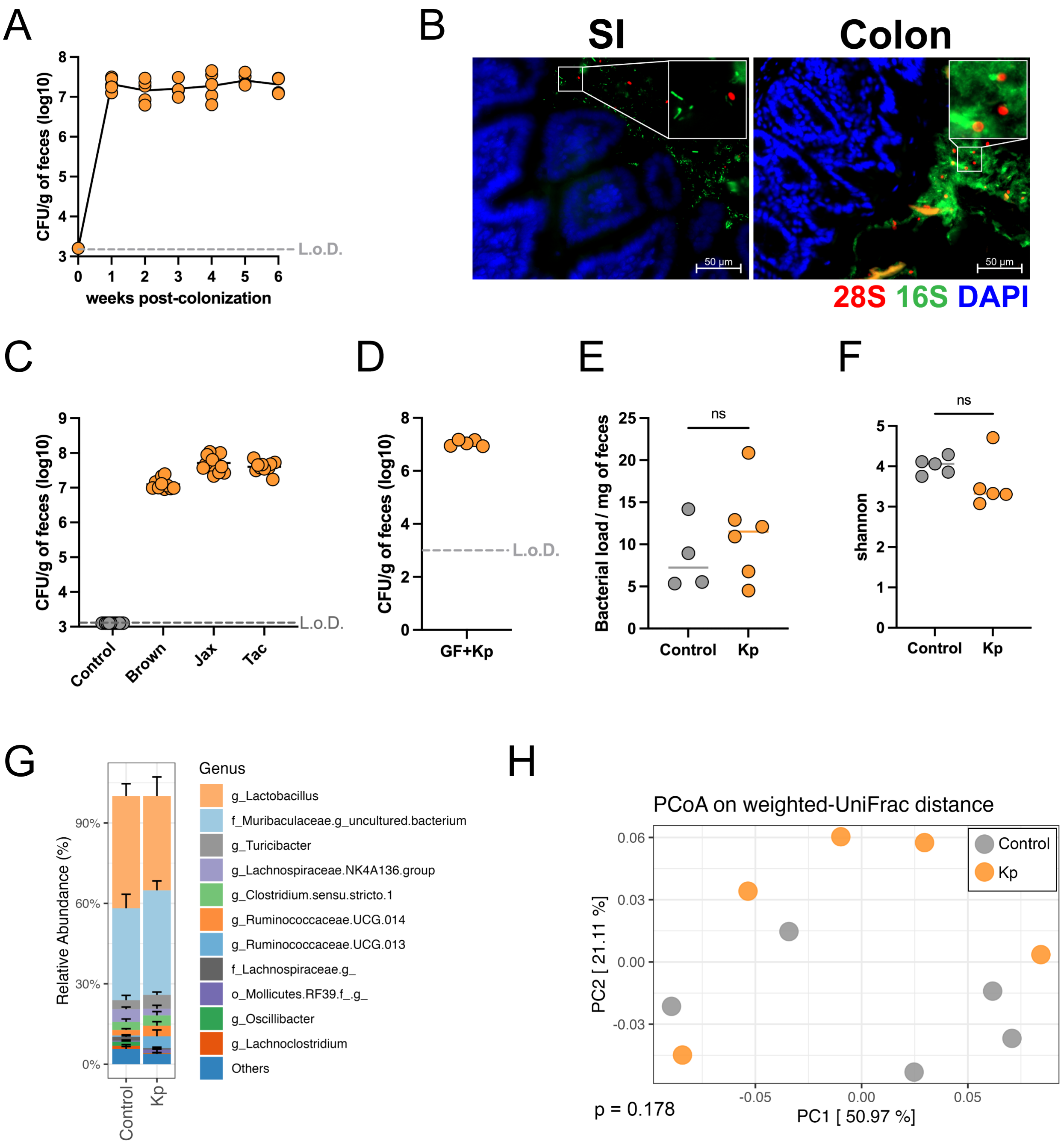


**Fig. S2.** ***K. pintolopesii* stably colonizes laboratory mice irrespective of gut bacteria and does not alter the bacterial communities.** (A) Fungal burden in the feces of mice after *K. pintolopesii* treatment (*n* = 5). (B) Representative images of *K. pintolopesii* colonization at 13 days post-colonization in the small intestine and colon.

Tissues were stained with DAPI (nuclei; blue), EUB338 probe (bacteria; green), and 28S probe (fungi; red). (C) Fungal burden in the feces of mice bred at Brown University (Brown; *n* = 10), Jackson Laboratory (Jax; *n* = 15), and Taconic Biosciences (Tac; *n* = 11) at 14 days post-colonization or without colonization (Control; *n* = 13) with *K. pintolopesii*. (D) Fungal burden in the feces of germ-free mice (GF; *n* = 5) at 14 days post-colonization with *K. pintolopesii*. (E) Bacterial load in the feces of the control and *K. pintolopesii*-colonized mice (Control, *n* = 4; Kp, *n* = 6). Bacterial load was expressed as a $2^{-Ct}$ value and normalized to the weight of the starting fecal material. (F-H) 16S rRNA gene sequencing was performed to explore fecal bacterial communities (Control, *n* = 5; Kp, *n* = 5). (F) Alpha diversity (Shannon) at 14 days post-colonization in the feces of control and *K. pintolopesii*-colonized mice. (G) Bar plot of the relative abundance of the bacterial genus from 16S rRNA gene sequencing of the feces. (H) PCoA plots based on the weighted-UniFrac distances at 14 days post-colonization in the feces. Significances were determined by PERMANOVA. Kp, *K. pintolopesii*-colonized mice.

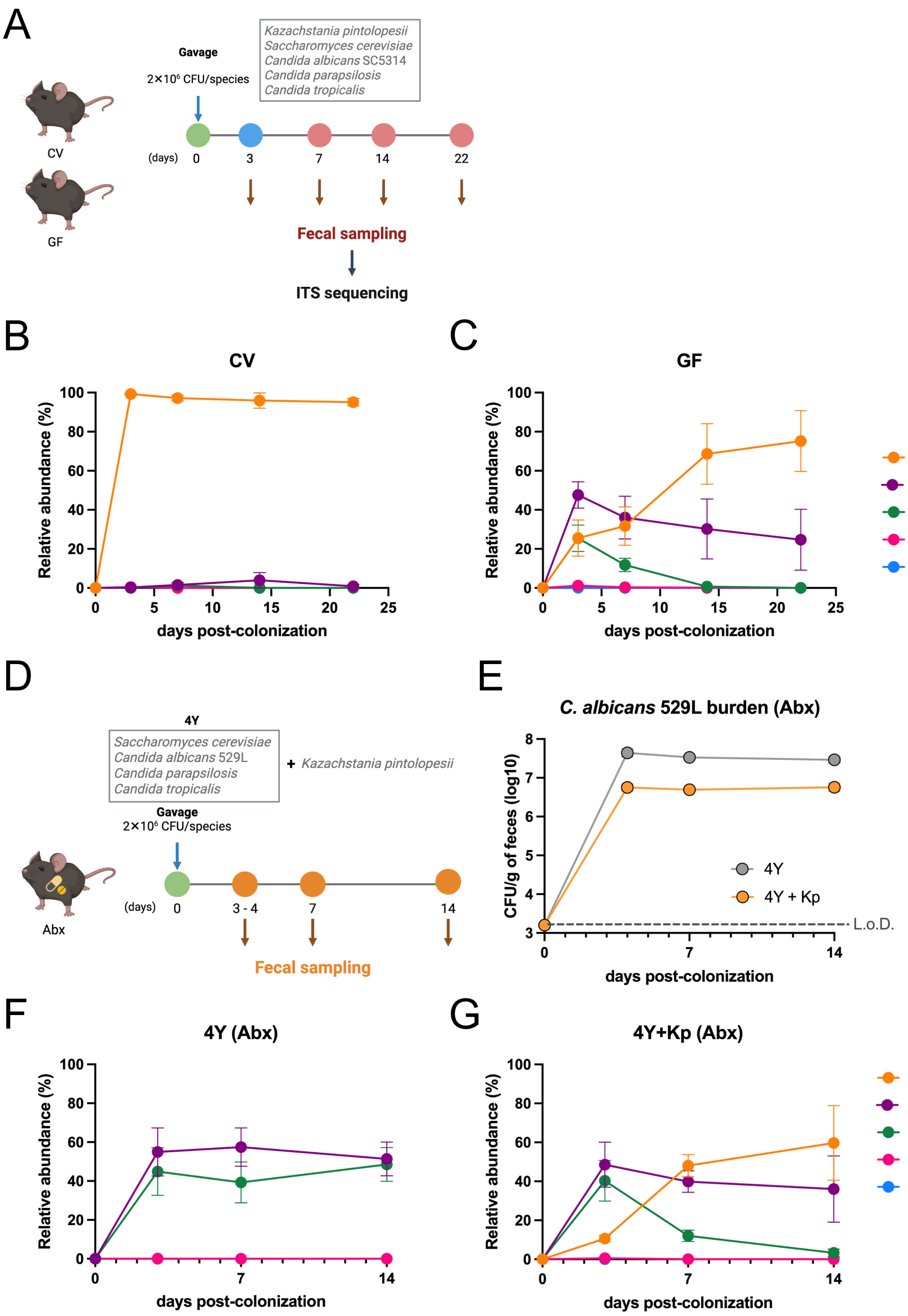


**Fig. S3. *K. pintolopesii* colonization limits the colonization of non-murine fungi.**

(A) Scheme of fungal competition experiment in the conventional and germ-free mice.

Line plot of relative abundance of fungal species from ITS-marker gene sequencing of the feces at different time points (days 3, 7, 14, and 22) in conventional (B) and germ-free mice (C) (*n* = 5 per group). (D) Scheme of fungal competition experiment in the antibiotic-treated mice (Abx). (E) Burden of the *C. albicans* 529L in Abx mice with (4Y + Kp) or without *K. pintolopesii* (4Y). Line plot of relative abundance of fungal species from ITS-marker gene sequencing of the feces at different time points (days 3, 7, and 14) in Abx mice without (F) or with *K. pintolopesii* (G) (*n* = 5 per group). Dots indicate mean ± SEM.

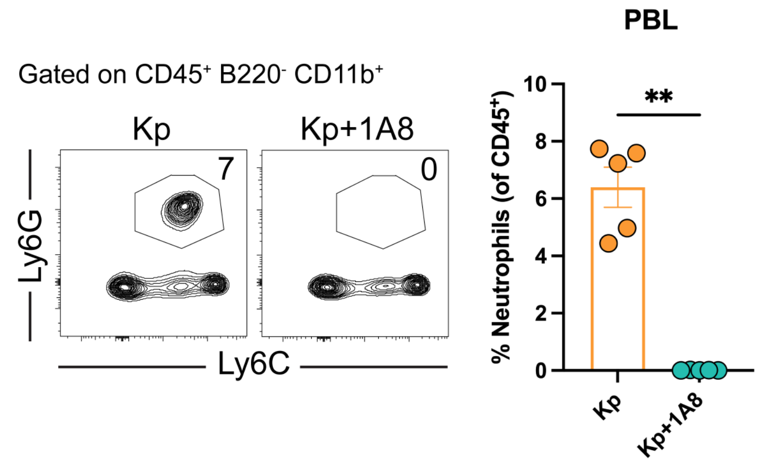


**Fig. S4. The evidence of neutrophil depletion after 1A8 antibody treatment.**

Representative flow cytometry plot and quantification of Ly6G$^{+}$ neutrophils after *K. pintolopesii* systemic infection (*n* = 5 per group). Bars indicate mean ± SEM. Significances were determined by the Mann-Whitney test. **p < 0.01. Kp, *K. pintolopesii*-colonized mice.

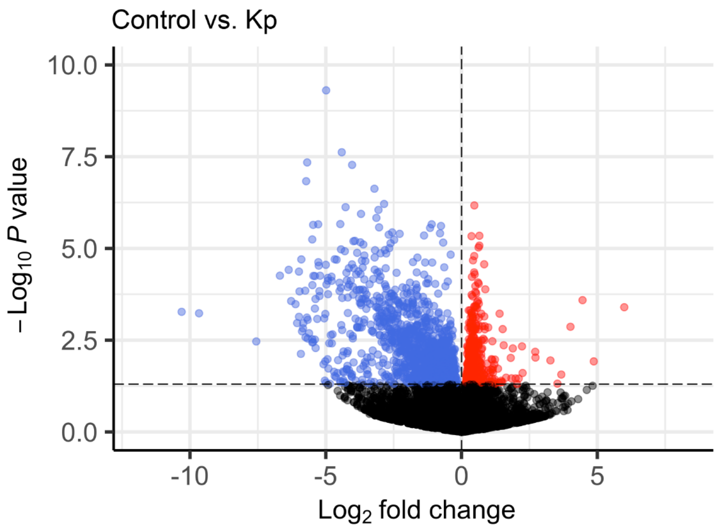


**Fig. S5. *K. pintolopesii* colonization drives Transcriptional changes in the small intestine.** Transcriptional profiles of the small intestines from control and *K. pintolopesii*-colonized mice were generated using bulk RNA sequencing ($n$ = 4 per group). Volcano plot of the transcriptional changes after *K. pintolopesii* colonization in the small intestine. Down-regulated and up-regulated genes after *K. pintolopesii* colonization were presented as blue and red dots, respectively. Kp, *K. pintolopesii*-colonized mice.

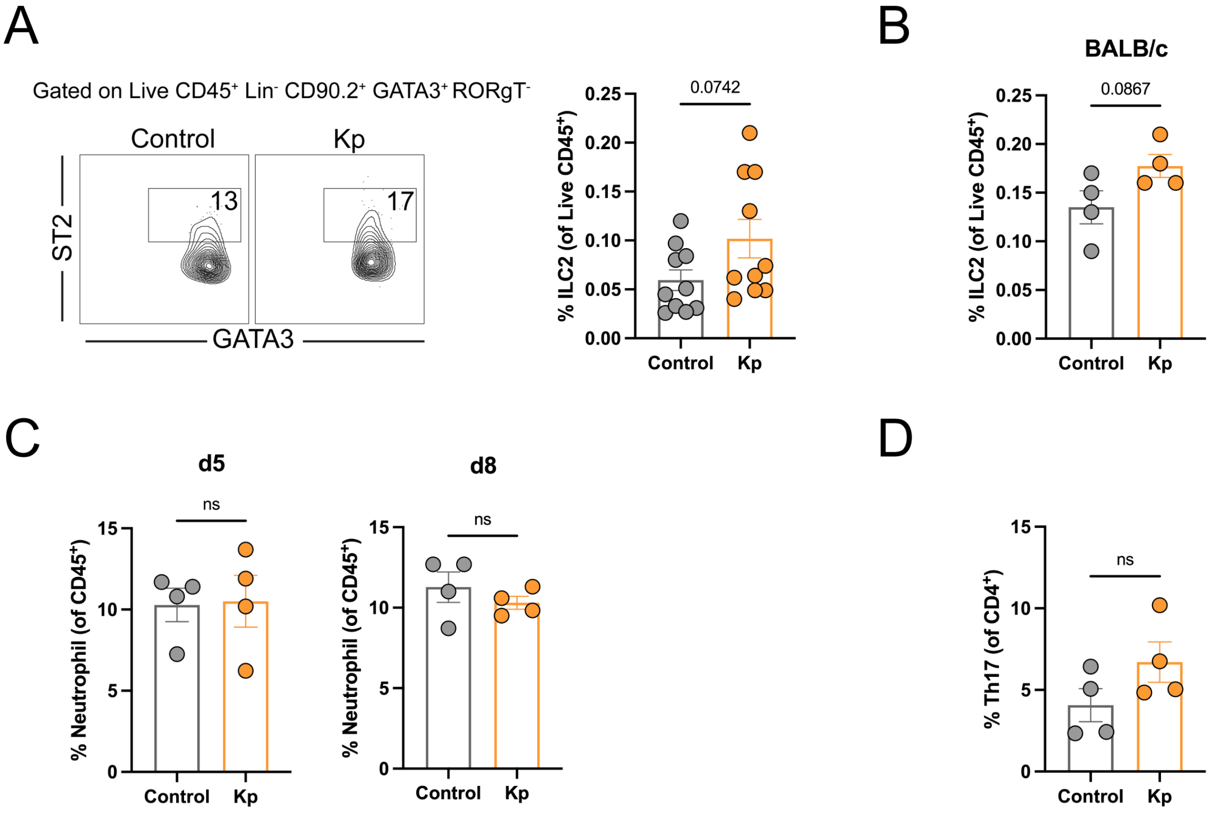


**Fig. S6. *K. pintolopesii* can colonize BALB/c mice without neutrophil and Th17 induction.** (A) Representative flow cytometry plots and frequency of ILC2 cells in the small intestinal lamina propria of control and *K. pintolopesii*-colonized mice at 14 days post-colonization (*n* = 10 per group). ILC2 populations were gated as live $CD45^{+}$ $Lin^{-}$ $CD90.2^{+}$ $GATA3^{+}$ $ROR\gamma T^{-}$ $ST2^{+}$. Lin: CD3ε, CD11b, Ly-6G/Ly-6C, CD19. (B) Frequency of ILC2 in the small intestinal lamina propria of control and *K. pintolopesii*-colonized BALB/c mice at 14 days post-colonization (*n* = 4 per group). (C) Frequency of circulating neutrophils in the peripheral blood at 5 and 8 days post-colonization of *K. pintolopesii* (*n* = 4 per group). Neutrophil populations were gated as $CD45^{+}$ $B220^{-}$ $CD11b^{+}$ $Ly6G^{+}$. (D) Frequency of Th17 cells in the small intestinal lamina propria at 14 days post-colonization of *K. pintolopesii* (*n* = 4 per group). Th17 populations were gated as live

$CD45^+$ $Lin^+$ $CD90.2^+$ $B220^-$ $CD4^+$ $CD8a^-$ $Foxp3^-$ $ROR\gamma T^+$. Lin: CD3ε, CD19. Bars indicate mean ± SEM. Kp, *K. pintolopesii*-colonized mice.
